# Subtypes of Bladder-Innervating Sensory Neurons revealed by Immuno-Fluorescence In Situ Hybridization

**DOI:** 10.64898/2026.09.03.749247

**Authors:** Xuejiao Sun, Guadalupe Manrique-Maldonado, Marcelo D. Carattino

## Abstract

Sensory neurons that innervate the urinary bladder are critical for triggering voiding and mediating pain perception under pathological conditions. Despite functional and immunohistochemical evidence indicating the existence of several bladder afferent populations, the subtypes of dorsal root ganglion (DRG) neurons that innervate this organ are poorly defined and specific markers to study them have not been described. Here, we combined retrograde tracing with the cholera toxin β subunit and immunofluorescence *in situ* hybridization with markers defined by transcriptomic analysis of C and Aδ afferents to classify the lumbosacral DRG neurons that innervate the urinary bladder. Four lumbosacral (L6-S2) bladder-innervating DRG neuron populations were identified: three *Calca*^+^ peptidergic populations that express *Adra2a*/*Th*, *Th*/*Ntrk2*, or *Ntrk2*, and one non-peptidergic population that expresses *Ntrk2*. Among the whole population of peptidergic lumbosacral sensory neurons, *Adra2a*^+^/*Th*^+^, *Th^+^*/*Ntrk2*^+^ and *Ntrk2*^+^ neurons were rare. Thus, our studies identified multiple populations of bladder-innervating DRG neurons, revealing unappreciated diversity among the sensory neuron subtypes present at the lumbosacral level that innervate visceral and somatic tissues.

## Introduction

The storage and release of urine depend on the function of sensory neurons that sense chemical and mechanical stimuli during the filling of the bladder, and communicate this information to the central micturition center in the brainstem [1, 2]. These neurons not only sense bladder distension but also function as nociceptors, encoding noxious stimuli. The cell bodies of the sensory neurons that innervate the urinary bladder reside in dorsal root ganglion (DRG) at lumbosacral (L6-S2; pelvic nerve afferents) and thoracolumbar (T13-L2; hypogastric nerve afferents) levels [1]. Sensory neurons are pseudo-unipolar with two axonal branches, a peripheral branch that transduces thermal, mechanical and chemical stimuli into electrical signals, and a central branch that relays these signals to the central nervous system for processing. Multiple subtypes of sensory neurons exist that encode a specific type or set of stimuli acting in the body, each with their own genetic identity, morphological characteristics, degree of myelination, and functional properties [3–6]. Yet, despite several decades of study, the molecular identity of the distinct sensory neuron subtypes that innervate the urinary bladder, their peripheral and central connections, and by extension the organization of the circuit underlying lower urinary tract control, remain not well understood.

Single-cell transcriptomic studies have identified at least 11 distinct DRG neuron subtypes in rodents and primates based on differential gene expression [3–6]. This includes five clusters of heavy myelinated (neurofilament rich, NF) proprioceptors and low threshold mechanoreceptors termed NF1, NF2, NF3, NF4 and NF5 that express *Ntrk2* and *Ntrk3*; three clusters of non-peptidergic (NP) nociceptors named NP1, NP2, and NP3, which express *Mrgprd*, *Mrgpra3* or *Sst*; peptidergic (PEP) non-myelinated C-fiber (PEP1) and lightly myelinated peptidergic fibers (PEP2) that express a variety of neuropeptides including calcitonin-gene related peptide alpha (*Calca*) and substance P (*Tac1*); and C-fiber low threshold mechanoreceptors that express *Th* [3, 4]. More recently, additional clusters of peptidergic DRG neurons expressing *Calca* were identified using single-cell (sc) RNA sequencing [7]. This expanded the number of subtypes of peptidergic nociceptors, although some of these clusters appear to overlap with previously described non-peptidergic neuronal subtypes (e.g., *Mrgpra3*).

While scRNA-seq and single-nucleus (sn) RNA-seq analyses have permitted the identification of molecularly distinct subtypes of somatosensory neurons [3–7], relatively few genes have been shown to be present in specific clusters and hence serve as true markers that could be used as surrogates for functional genotyping across multiple tissues. Validation in transcriptomic studies is typically limited to a few genes, with cross-interrogation of markers rarely performed. For the urinary bladder and other visceral organs, it is currently unknown whether they are innervated by unique sensory neuron subtypes or, alternatively, by previously defined sensory neuron populations. In the case of colonic DRG neurons, scRNA-seq revealed molecular types distinct from those reported in other general transcriptomic studies [8, 9], suggesting the existence of subsets of sensory neurons dedicated to innervation of visceral organs. Thus, the objectives of this study were twofold: to define molecular markers for the DRG neuron subtypes that innervate the urinary bladder, and to assess whether the neuronal populations that innervate this organ exhibit different molecular profiles to those innervating other tissues. To accomplish this, we combined retrograde labeling with cholera toxin β subunit and immuno-fluorescence *in situ* hybridization.

## Materials and methods

### Reagents

All chemicals were purchased from Sigma-Aldrich (St. Louis, MO), unless otherwise specified.

### Mice

Experimental procedures were approved by the University of Pittsburgh Institutional Animal Care and Use Committee. Female C57BL/6J mice were purchased from the Jackson Laboratory and housed in standard cages at the University of Pittsburgh under 12h light/12h dark cycles with free access to food and water. All experimental mice were 3-6 month-old virgin females and group housed after weaning up to 5 per cage. Animals were euthanized by CO_2_ inhalation, followed by a thoracotomy.

### Retrograde labeling of bladder sensory neurons

Bladder-innervating sensory neurons were labeled with cholera toxin β (CTb) subunit (Millipore-Sigma, Cat. N° C9903-1mg). CTb was dissolved in sterile saline to a final concentration of 0.5%. Mice were anesthetized with isoflurane and an abdominal incision was made to expose the urinary bladder. CTb was injected into the ventral and dorsal aspects of the bladder dome at 3 to 4 sites (2 μl per site) with a Hamilton^TM^ 600 Microliter syringe (Hamilton, Cat. N°. 763301) suited with a 33G needle (Hamilton, Cat. N°. 7803-15). The abdominal incision was closed in layers with 5.0 PDO absorbable monofilament surgical sutures (AD Surgical, Cat. N°. S-D518R13). Ketoprofen (5 mg/kg) was administered subcutaneously to alleviate pain (5 mg/kg, Zoetis, Ketofen) and ampicillin (100 mg/kg, Eugia US LLC, Cat. No. NDC 55150-113-10) to prevent infections.

### Tissue preparation and sectioning

Lumbosacral (L6-S2) DRG were harvested 7 days after the surgical procedure. Mice were euthanized by CO_2_ inhalation following by a thoracotomy. Blood was removed by intracardiac perfusion with sterile saline. For fluorescence *in situ* hybridization (FISH) and immuno-FISH, L6-S2 DRG were collected, placed into a 35 mm Petri dish filled with cold Neurobasal A media (ThermoFisher Scientific, Cat. No. 10888022). Then, DRG were transferred to a 35 mm Petri dish filled with Optimal Cutting Temperature embedding medium (OCT; Fisher Scientific, Cat. N°. 23-730-571) and subsequently were placed in the bottom of a 10 x 10 x 5 mm Tissue-Tek Cryomold (Sakura Finetek USA Inc, Cat. N°. 427971) filled with OCT. Tissues were frozen by placing the blocks in a -80°C freezer.

### Tissue processing

DRG sections (12 μm) were cut with a Leica CM1950 cryostat (chamber temperature of -20°C and a knife temperature of -18°C), collected on SuperfrostTM Plus microscope slides (ThermoFisher Scientific, Cat. N°. 12-550-15). Slides containing sectioned tissues were allowed to “dry” in the cryostat chamber for 30 min, before long-term storage at - 80°C.

### Fluorescence in situ hybridization (FISH)

The RNAscope^TM^ Multiplex Fluorescent v2 kit (ACD, Cat. N°. 323100) was used to examine gene expression in DRG and implemented according to ACD guidelines with some modifications. Probes for genes of interest are listed in Table I. The 3-plex negative control probe against Bacillus subtilis dapB (ACD, Cat. No. 320871) was used to detect non-specific signals. Slides containing sectioned tissue were removed from the -80 °C freezer and immediately fixed for 15 min at 4 °C with neutral buffered formalin (29 mM NaH_2_PO_4_•H_2_O, 45.8 mM Na_2_HPO_4_, and 4.0% (v/v) paraformaldehyde). Then, slides were washed with PBS and subsequently incubated with Protease IV for 5 min at room temperature. Probes were developed using TSA Vivid^TM^ Fluorophore 520 (TocrisBiosciences, Cat. N°. 7523), TSA Vivid^TM^ Fluorophore 570 (Tocris Biosciences, Cat. N°. 7526). Nuclei were counterstained with DAPI. Tissues sections were mounted under borosilicate coverslips (ThermoFisher, Cat. No. 12541019 or Cat. No. 12541024) using ProLong Gold Antifade mounting medium (ThermoFisher, Cat. No. P36934) and cured overnight at room temperature in the dark. The slides were then stored at 4 °C.

**Table I.** RNAscope probes used in this study.

| RNA probe | Cat. No | Channel |
| --- | --- | --- |
| *Mm-Calca* | 578771 | C2, C3 |
| *Mm-Adcyap1* | 405911 | C1, C3 |
| *Mm-Bmpr1b* | 533941 | C1 |
| *Mm-Mrgprd* | 417921 | C1, C3 |
| *Mm-Mrgpra3* | 548161 | C2 |
| *Mm-Adra2a* | 425341 | C3 |
| *Mm-Th* | 317621 | C1, C3 |
| *Mm-SST* | 482691 | C1 |
| *Mm-Ntrk2* | 423611 | C2, C3 |
| *Mm-Adra2a* | 425341 | C3 |
| *Mm-Piezo2* | 400191 | C3 |

### Immuno-FISH

Slides containing sectioned tissue were removed from the -80 °C freezer and immediately fixed for 15 min at 4 °C with neutral buffered formalin. Then, sections were incubated overnight with primary antibodies diluted in codetection antibody diluent (ACD, Cat. No. 323160) at 4 °C. Primary antibodies used were rabbit anti-Cholera Toxin (Novus, Cat. No. 100-63067, dilution 1:100), goat anti-CTb (Sigma-Millipore, Cat. No. 227040, dilution 1:1000) and rabbit anti-TH (AB152, Sigma-Millipore, dilution 1:100). After primary antibody incubation, slides were incubated in neutral buffered formalin for 30 min at room temperature, and then with Protease IV for 5 min at room temperature. Probes were developed as indicated above. Following the last treatment with horseradish peroxidase blocker, samples were incubated for 30 min at room temperature with secondary antibodies. Secondary antibodies used were donkey anti-rabbit conjugated to AlexaFluor^TM^ 647 (Jackson Labs Antibodies, Cat. No. 711-605-152, dilution 1:100) and donkey anti-goat conjugated to AlexaFluor^TM^ 568 (Jackson Labs Antibodies, Cat. No. 705-575-147, dilution 1:100). Nuclei were counterstained with DAPI and sections were mounted with ProLong Gold antifade mountant, as indicated above.

### Image Capture

A Leica Microsystems SP8 Stellaris confocal microscope outfitted with a 405-laser diode and a white light laser and a Leica HCX PL APO 20x, 0.75 numerical aperture dry objective was used to image labeled DRG sections. 8-bit images were collected using 2-line averages combined with 2 frame averages. Cross-talk between channels was prevented by use of spectral detection coupled with sequential scanning. Stacks of images (1,024 x 1,024, 8-bit) were collected using system-optimized parameters for the z-axis. Images were processed using the 3D visualization package of Leica LASX software and exported as TIFF files. Final figures were assembled in Adobe Illustrator version 29.6.1 (Adobe).

### Image analysis and quantification

All images analysis was performed with ImageJ/FIJI. Only neurons with visible nucleus were analyzed. To quantify the percentage of DRG neurons positive for a specific marker in FISH studies, the nucleus of individual neurons was identified using DAPI staining, and the polygon tool was used to draw regions of interest (ROI) around the nucleus. For immuno-FISH studies, ROIs were draw around neurons based on the CTb signal. The *Ntrk2* is expressed in sensory neurons and satellite glial cells. To avoid false positives, *Ntrk2* signal was assessed exclusively within the nucleus of CTb-labeled neurons. After drawing the ROI, neurons were numbered in sequential order. Neurons displaying five or more puncta above the level observed in negative control slides were considered positive for that marker. The number of neurons positive for one or both markers tested was computed manually for each section. For whole DRG neuron analysis, data are from DRG sections of at least three animals, and each data point corresponds to the percentage of positive neurons for the specific marker in an individual section. For CTb-labeled neurons, the percentage of positive neurons for a given marker was calculated from at least 2 tissue blocks containing DRG from 2-3 mice injected with CTb. The number of tissue sections analyzed is indicated in the figure legends.

### Statistical Analysis

Data are presented as mean ± SEM (*n*), where n equals the number of independent measurements.

## RESULTS

### Selection of DRG neuron markers

To characterize the sensory neuron subtypes innervating the urinary bladder, we selected molecular markers that were previously shown to label transcriptionally defined clusters of unmyelinated and lightly myelinated DRG neuron including: *Adra2a* (CGRP-γ;PEP1), *Bmpr1b* (CGRP-η;PEP2), *Mgrpra3* (CGRP-θ_1_;NP2), *Mgrprd* (NP1), *Ntrk2* (Aδ-LTMR), *Sst* (NP3) and *Th* (TH; C-LTMRs) [3, 4, 7]. *Calca*, which encodes for the neuropeptide calcitonin gene-related peptide (CGRP), was used as a general marker of peptidergic neurons. Earlier work has established that heavily myelinated neurons with conduction velocities in the Aα and Aβ range do not innervate the urinary bladder and therefore were not targeted [10–19].

### Validation of markers for lumbosacral DRG neuron

We initially used fluorescence *in situ* hybridization (FISH) to evaluate the specificity of the selected markers to label lumbosacral (L6-S2) sensory neuron subtypes. These markers are thought to label unique DRG neuron subtypes in mouse [7, 20]. Fresh frozen L6-S2 DRG sections were incubated with various combinations of RNAScope^TM^ probes; with each probe tested against every other probe. For each combination of markers, the percentage of single or double-labeled DRG neurons was calculated (Fig. 1 and 2). Consistent with previous studies [7], we confirmed that *Mgrpra3*, *Mgrprd*, and *Sst* label unique DRG neuron subsets at the lumbosacral level. *Mgrpra3* is expressed in the CGRP-θ_1_ cluster, a population of CGRP^+^ C-fibers. Some degree of overlap between markers was noted with certain probe combinations (Fig. 2). For instance, the majority of *Adra2a*^+^ neurons were peptidergic, expressing *Calca* (85.7±5.3%). A fraction of peptidergic *Adra2a*^+^ neurons also co-express *Bmpr1b* (28.3±1.2%) and *Th* (34.3±9.3 %) (Fig. 2). Likewise, two clusters of *Ntrk2*^+^ sensory neurons were identified: one that was negative for all molecular markers tested, and another that expresses *Bmpr1b* (32.9±6.6 %). Of the *Th*^+^ lumbosacral sensory neurons, 20.2±7.6% express *Adra2a* and 13.3±7.3% express *Bmpr1b*. In summary, FISH studies indicate that while some markers reliably define discrete neuronal populations, others reveal previously underappreciated molecular heterogeneity within DRG neurons, underscoring the need for combinatorial marker strategies to accurately classify bladder-innervating DRG neuron subtypes.

**Figure 1.**
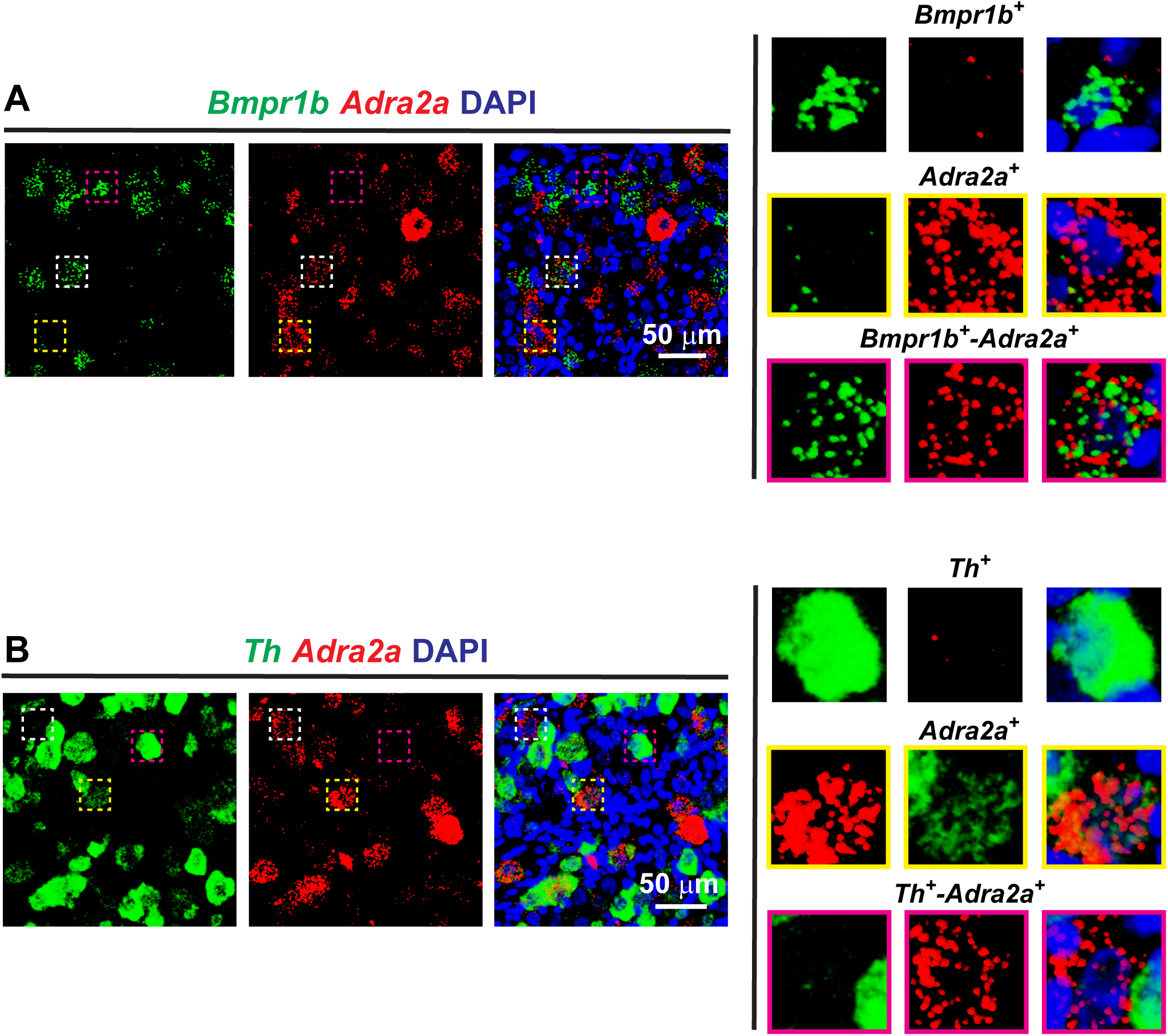
Multiple sensory neuron population at the lumbosacral level express *Bmpr1b, Adra2a and Th*. FISH was performed in fresh frozen sections of lumbosacral dorsal root ganglia DRG. (**A**) Example of confocal images of DRG labeled with probes for *Adra2a* and *Bmpr1b*. (**B**) Example of confocal images of DRG labeled with probes for *Adra2a* and *Th*. Inset, 4-fold magnification of boxed sensory neurons.

**Figure 2.**
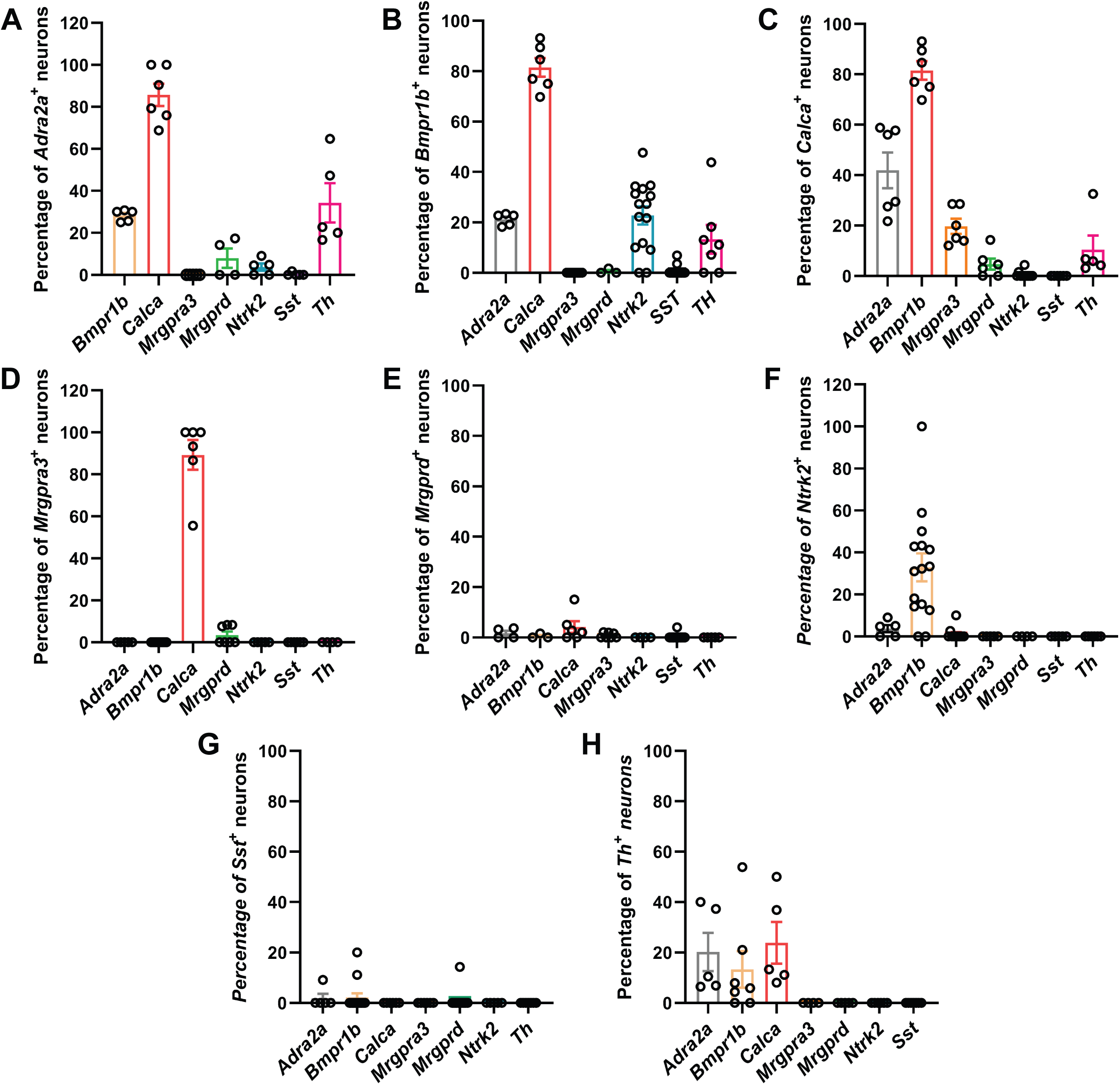
Validation of markers for lumbosacral DRG neuron. FISH was used to evaluate the specificity of the selected markers to label lumbosacral (L6-S2) sensory neurons. Fresh frozen DRG sections were incubated with various combinations of probes. Every probe was tested against all other probes. (**A**-**H**) Percentage of *Adra2a*^+^ (**A**), *Bmpr1b*^+^ (**B**), *Calca*^+^ (**C**), *Mrgpra3*^+^ (**D**), *Mrgprd*^+^ (**E**), *Ntrk2*^+^ (**F**), *Sst*^+^ (**G**) and *Th*^+^ (**H**) neurons expressing each marker. Dots indicate the number of DRG sections analyzed for each combination. Data are shown as the mean ± SEM.

### Bladder-innervating sensory neuron subtypes

To define molecular markers for the sensory neurons that innervate the urinary bladder, we combined retrograde labeling with cholera toxin β subunit (CTb) and immuno-FISH. A total of 3088 CTb labeled DRG neurons were analyzed. Consistent with previous studies showing that a large fraction of bladder afferents expresses the neuropeptide CGRP [2], we found that approximately 80.9±1.7% of the retrogradely labeled neurons express mRNA encoding for *Calca*. We observed substantial overlap between *Bmpr1b* expression and that of other markers in CTb-labeled neurons; therefore, this marker was excluded from further analysis. 49.6±2.1% of the bladder-innervating sensory neurons express *Adra2a* (Fig. 3). Most *Adra2a*^+^ sensory neurons labeled with CTb, as expected express *Calca*, and also *Th* (Fig. 3). In contrast, a relatively small percentage of unlabeled lumbosacral *Adra2a*^+^ sensory neurons express *Th* (34.3±9.3 %) (Fig. 2). *Ntrk2*, a marker of myelinated neurons [3, 4], is expressed in approximately 40.8±2.1 % of the bladder-innervating DRG neurons, and 28.3±0.5 % of them are also positive for *Calca* (Fig. 4). Of the peptidergic *Ntrk2*^+^ bladder-innervating DRG neurons, 10.7±3.9 % express *Th* (Fig. 4). Bladder-innervating sensory neurons do not express *Mgrpra3*, *Mgrprd*, or *Sst* (Fig. 5). In summary, *Adra2a*^+^/*Th*^+^/*Calca*^+^ constitute a well-defined population that accounts for almost 50 % of the bladder-innervating neurons. The non-peptidergic *Ntrk2* population is defined by the absence of *Calca*, and it resembles a population of previously identified Aδ-LTMR [20]. The other two peptidergic populations express *Ntrk2* and are defined by the presence or absence of *Th expression*. Taken as a whole, our data indicate that four major DRG neuron subtypes innervate the urinary bladder: *Adra2a*^+^/*Th*^+^/*Calca*^+^ (PEP1); Ntrk2^+^/*Th*^+^/*Calca*^+^ (PEP2); *Ntrk2^+^*/*Calca*^+^ (PEP3); and *Ntrk2*^+^ (NP1) (Fig. 3-4).

**Figure 3.**
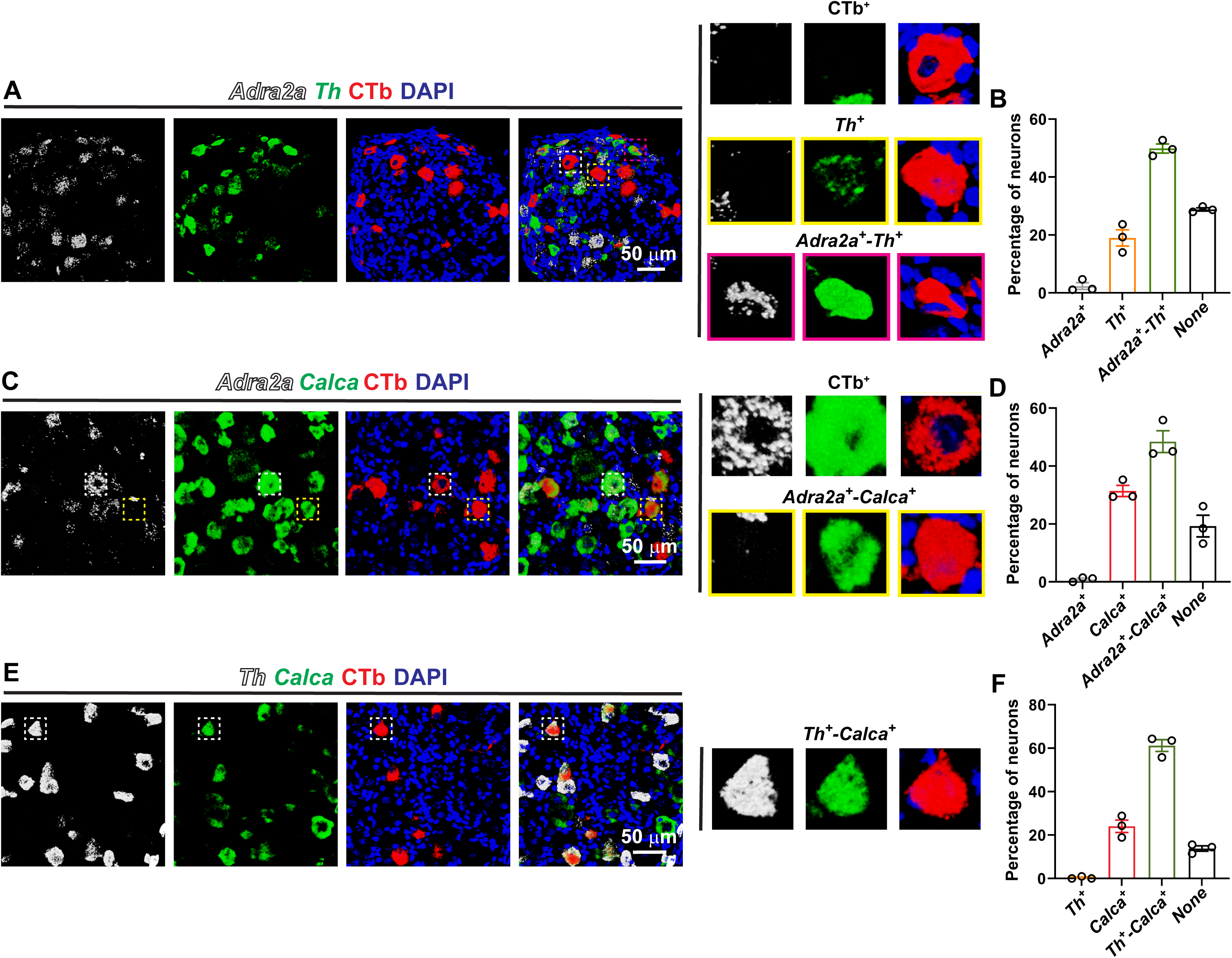
Peptidergic *Adra2a*^+^-*Th*^+^ *neurons* innervate the urinary bladder. Immuno-FISH was performed in fresh frozen sections of DRG (L6-S2) harvested from mice injected into the bladder wall with cholera toxin β subunit (CTb). A rabbit antibody anti-CTb and a secondary donkey anti-rabbit conjugated with AlexaFluor^TM^ 647 were used to identify bladder sensory neurons (red). (**A**) Example of confocal images of DRG labeled with probes for *Adra2a* and *Th*. Inset, 4-fold magnification of CTb-labeled neurons. Note that most *Adra2a*^+^ neurons express *Th*. (**B**) Percent of CTb-labeled cell bodies that express *Adra2a, Th*, or both. Data from N=35 images from 6 mice. (**C**) Example of confocal images of DRG labeled with probes for *Adra2a* and *Calca*. Inset, 4-fold magnification of CTb-labeled neurons. Note that most *Adra2a*^+^ neurons express *Calca*. (**D**) Percent of CTb-labeled cell bodies that express *Adra2a, Calca*, or both. Data from N=30 images from 6 mice. (**E**) Example of confocal images of DRG labeled with probes for *Th* and *Calca*. Inset, 4-fold magnification of CTb-labeled neurons. Note that most *Th*^+^ neurons express *Calca*. (**F**) Percent of CTb-labeled cell bodies that express *Th, Calca*, or both. Data from N=32 images from 6 mice. Data are shown as the mean ± SEM.

**Figure 4.**
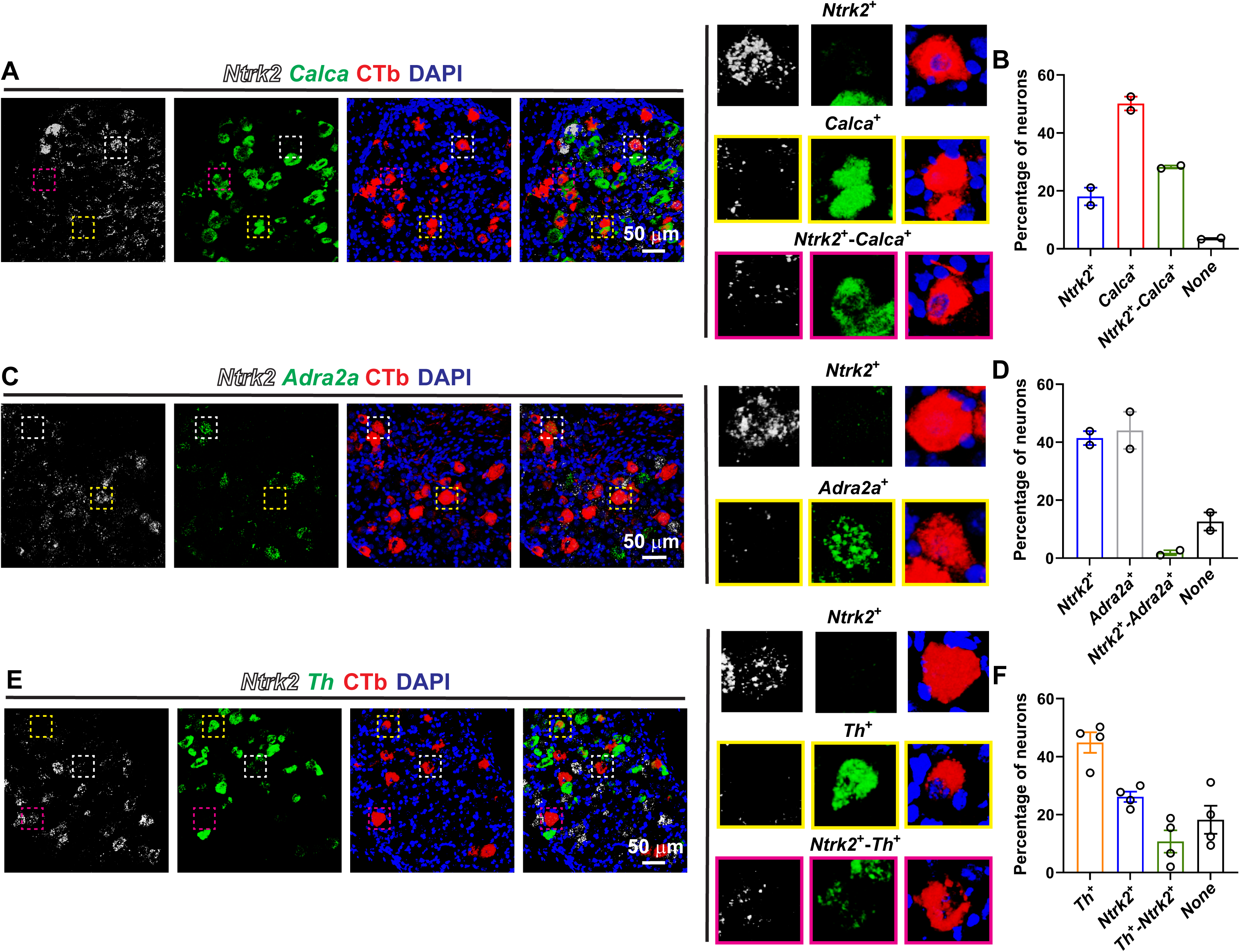
Several *Ntrk2*^+^ sensory *neuron populations* innervate the urinary bladder. Immuno-FISH was performed in fresh frozen sections of DRG (L6-S2) harvested from mice injected into the bladder wall with cholera toxin β subunit (CTb). A rabbit antibody anti-CTb and a secondary donkey anti-rabbit conjugated with AlexaFluor^TM^ 647 were used to identify bladder sensory neurons (red). (**A**) Example of confocal images of DRG labeled with probes for *Ntrk2* and *Calca*. Inset, 4-fold magnification of CTb-labeled neurons. (**B**) Percent of CTb-labeled cell bodies that express *Ntrk2, Calca*, or both. Data from N=18 images from 4 mice. (**C**) Example of confocal images of DRG labeled with probes for *Ntrk2* and *Adra2a*. Inset, 4-fold magnification of CTb-labeled neurons. (**D**) Percent of CTb-labeled cell bodies that express *Ntrk2, Adra2a*, or both. Data from N=22 images from 4 mice. (**E**) Example of confocal images of DRG labeled with probes for *Ntrk2* and *Th*. Inset, 4-fold magnification of CTb-labeled neurons. (**F**) Percent of CTb-labeled cell bodies that express *Ntrk2, Th*, or both. Data from N=18 images from 4 mice. Data are shown as the mean ± SEM.

**Figure 5.**
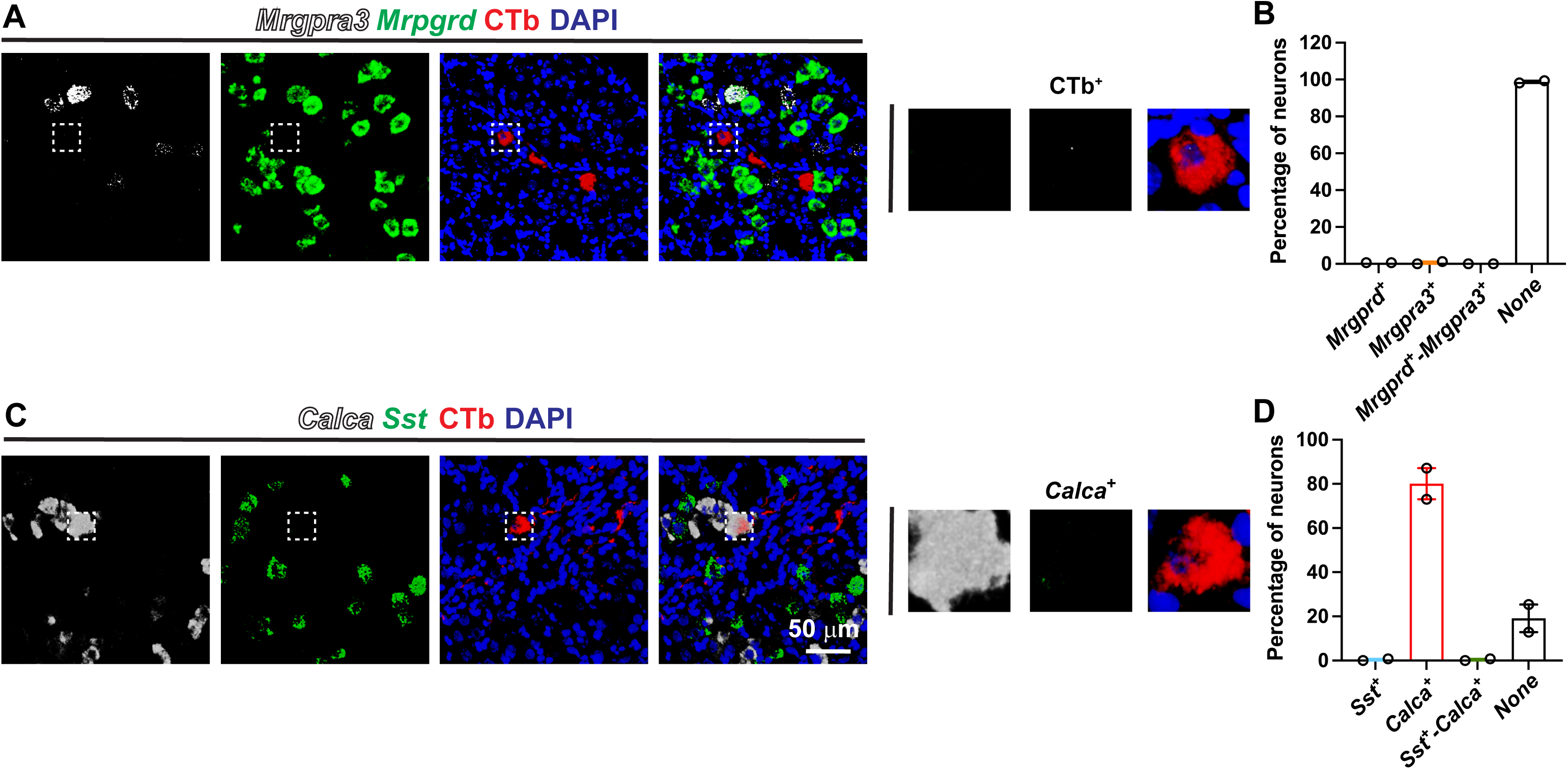
*Mrpgrd3*, *Mrpgrd* and *Sst* sensory neurons do not innervate the urinary bladder. Immuno-FISH was performed in fresh frozen sections of DRG (L6-S2) harvested from mice injected into the bladder wall with cholera toxin β subunit (CTb). A rabbit antibody anti-CTb and a secondary donkey anti-goat conjugated with AlexaFluor^TM^ 647 were used to identify bladder sensory neurons (red). (**A**) Example of confocal images of DRG labeled with probes for *Mrpgra3* and *Mrpgrd*. Inset, 4-fold magnification of CTb-labeled neurons. (**B**) Percent of CTb-labeled cell bodies that express *Mrpgra3* or *Mrpgrd*. Data from N=22 images from 4 mice. (**C**) Example of confocal images of DRG labeled with probes for *Sst* and *Calca*. Inset, 4-fold magnification of CTb-labeled neurons. (**D**) Percent of CTb-labeled cell bodies that express *Sst, Calca*, or both. Data from N=21 images from 4 mice.

### Validation of neuronal markers at protein level

Our studies identified four distinct subtypes of bladder-innervating neurons, each defined by a unique combination of marker genes. To confirm these transcriptionally defined subtypes at protein level, we performed immuno-FISH with antibodies against TH and CTb and probes for the main markers identified above. We did not find commercially suitable antibodies for immuno-FISH against ADRA2A or NTRK2, and therefore, we used mRNA probes instead. Bladder-innervating sensory neurons were retrogradely labeled with CTb as described previously. Consistent with the FISH labeling, almost all retrogradely labeled DRG neurons that express TH were peptidergic (*Calca*^+^) (Fig. 6A-B). While *Adra2a* was predominantly expressed in TH^+^ CTb-labeled neurons, not all TH-positive neurons co-expressed *Adra2a* (Fig. 6C-D). A small but significant number of *Adra2a*^+^ neurons (6.3±2.5) did not express TH; a representative example is shown in Fig. 6C. Lastly, we confirm the existence of two major populations of *Ntrk2*^+^ bladder-innervating DRG neurons, distinguished by the presence or absence of TH expression (Fig. 6E-F). Thus, orthogonal validation using immuno-FISH and an antibody against TH supports the distinction between *Th*^+^ and *Th*-neg populations, further substantiating the existence of four distinct DRG neuron populations innervating the urinary bladder.

**Figure 6.**
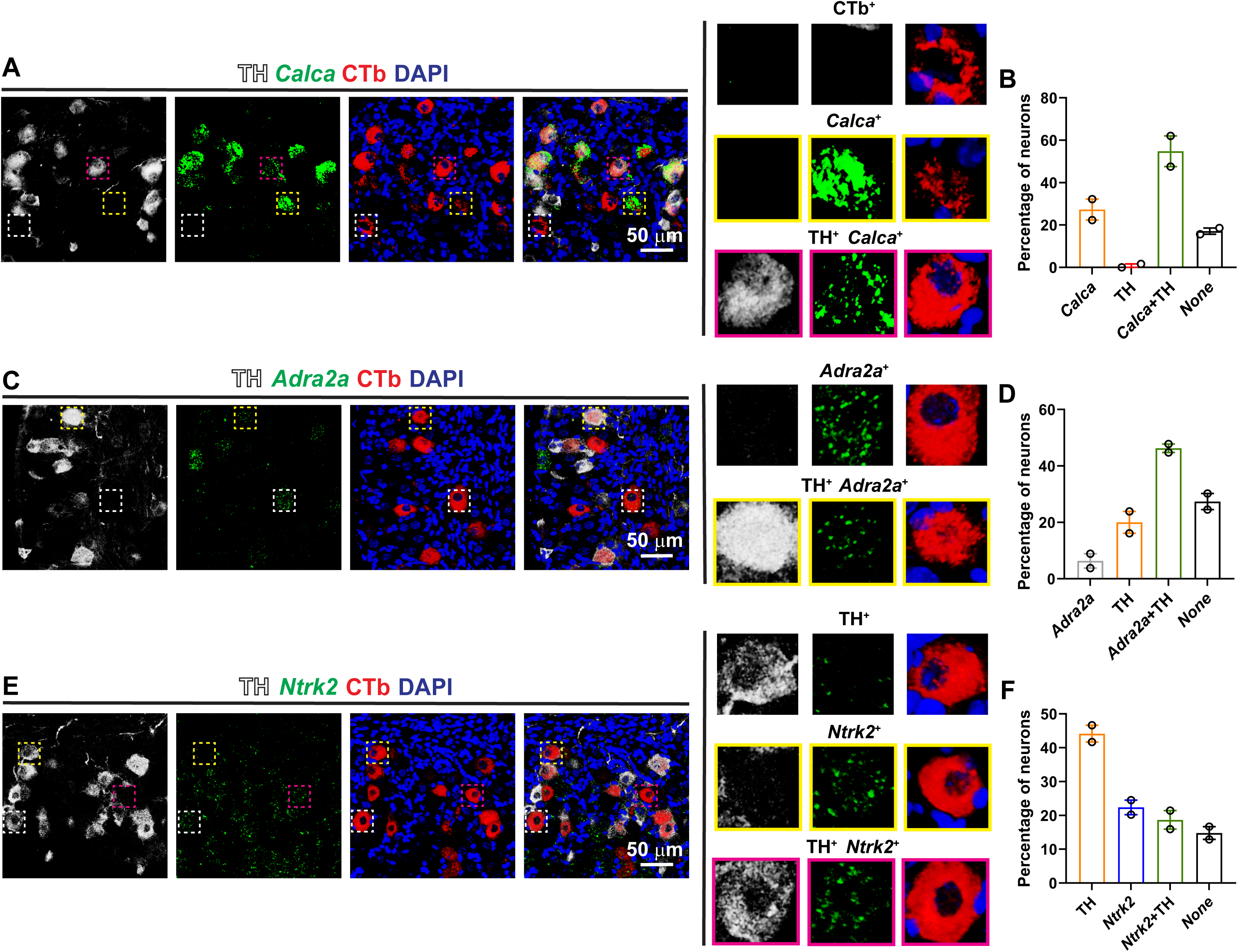
Bladder-innervating DRG neuron subtypes revealed by immuno-FISH. Mice were injected into the bladder wall with cholera toxin β subunit (CTb). Immuno-FISH was performed in fresh frozen sections of DRG (L6-S2). A goat antibody anti-CTb and a secondary donkey anti-goat conjugated with AlexaFluor^TM^ 568 were used to identify bladder sensory neurons (red). A rabbit antibody and a secondary donkey anti-rabbit conjugated with AlexaFluor^TM^ 647 were used to detect TH (white). (**A**) Example of confocal images of DRG labeled with antibodies against CTb and TH and a probe for *Calca*. Inset, 4-fold magnification of CTb-labeled neurons. Note that most of the neurons that express TH are peptidergic. (**B**) Percent of CTb-labeled cell bodies that express *Calca,* TH, or both. Data from N=15 images from 4 mice. (**C**) Example of confocal images of DRG labeled with antibodies against CTb and TH and a probe for *Adra2a*. Inset, 4-fold magnification of CTb-labeled neurons. Note that a small percentage of *Adra2a*^+^ neurons is negative for TH. (**D**) Percent of CTb-labeled cell bodies that express *Adra2a,* TH, or both. Data from N=15 images from 4 mice. (**E**) Example of confocal images of DRG labeled with probes for TH and *Ntrk2*. Inset, 4-fold magnification of CTb-labeled neurons. (**F**) Percent of CTb-labeled cell bodies that express TH*, Ntrk2*, or both. Data from N=16 images from 4 mice. Data are shown as the mean ± SEM.

### Piezo2 is expressed in Ntrk2-positive sensory neurons

To further assess the specificity of the identified molecular markers to label distinctive DRG neuron subtypes, we investigated the expression of mRNA encoding the mechanically activated ion channel *Piezo2*. We reason that if the defined markers label unique populations of bladder-innervating sensory neurons, *Piezo2* expression should be limited to a few subtypes. As shown in Fig. 7, *Piezo2* is found primarily in bladder-innervating sensory neurons that express *Ntrk2* including *Calca*^+^ and *Calca*-neg. In summary, our studies indicate that *Piezo2* expression is confined to *Ntrk2*^+^ bladder-innervating sensory neurons, meaning PIEZO2-mediated mechanosensation in the bladder is likely molecularly restricted to defined subsets of afferents.

**Figure 7.**
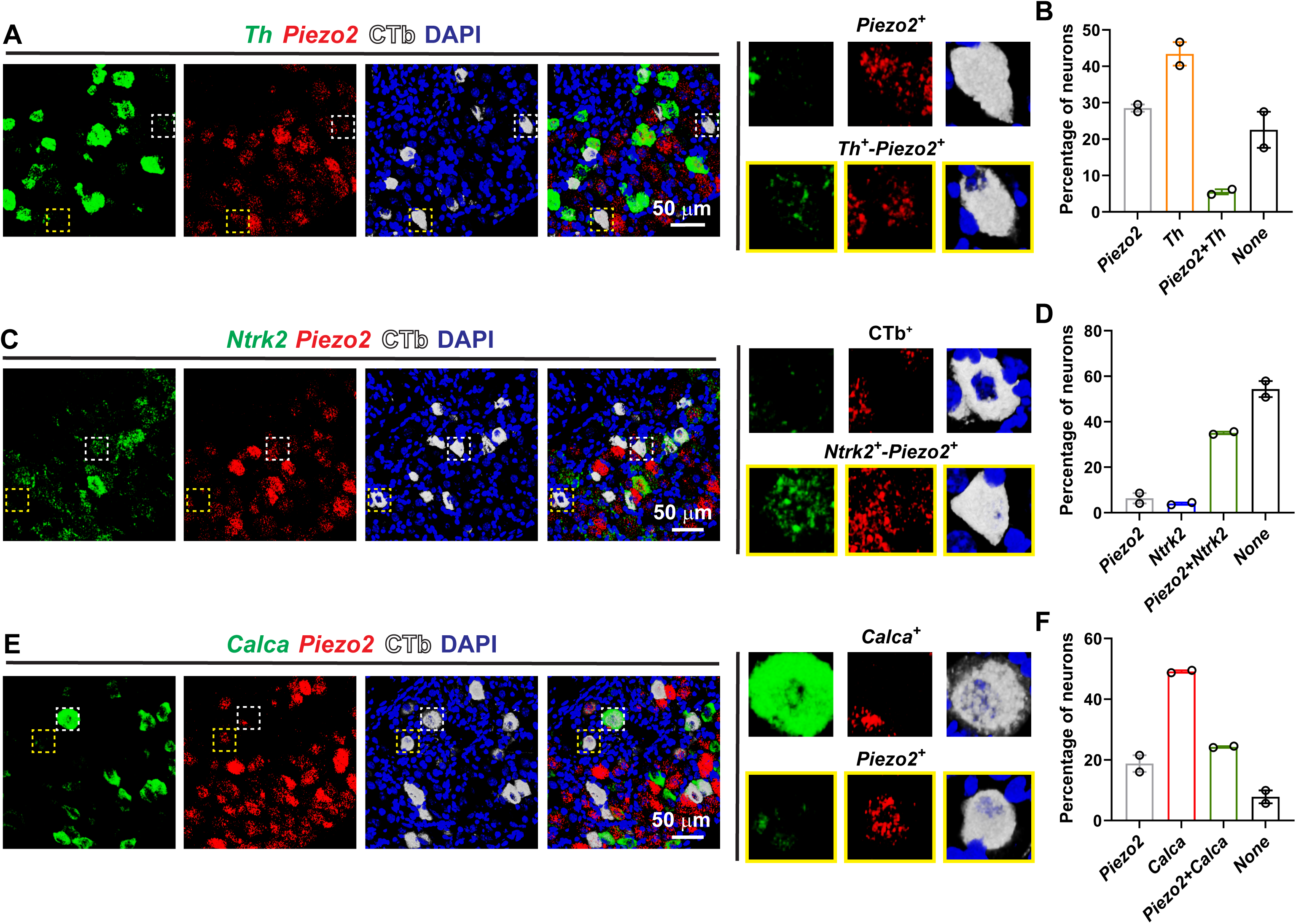
*Piezo2 is expressed in Ntrk2*^+^ bladder-innervating sensory neurons. Immuno-FISH was performed in fresh frozen sections of DRG (L6-S2) collected from mice injected into the bladder wall with cholera toxin β subunit (CTb). A rabbit antibody anti-CTb and a secondary donkey anti-rabbit conjugated with AlexaFluor^TM^ 647 were used to identify bladder sensory neurons (white). (**A**) Example of confocal images of DRG labeled with probes for *Piezo2* and *Th*. Inset, 4-fold magnification of CTb-labeled neurons. (**B**) Percent of CTb-labeled cell bodies that express *Piezo2*, *Th*, or both. Data from N=21 images from 4 mice. (**C**) Example of confocal images of DRG labeled with probes for *Piezo2* and *Ntrk2*. Inset, 4-fold magnification of CTb-labeled neurons. Note that most *Ntrk2*^+^ neurons express *Piezo2*. (**D**) Percent of CTb-labeled cell bodies that express *Piezo*2*, Ntrk2*, or both. Data from N=21 images from 4 mice. (**E**) Example of confocal images of DRG labeled with probes for *Piezo2* and *Calca*. Inset, 4-fold magnification of CTb-labeled neurons. (**F**) Quantification of the percent of CTb-labeled cell bodies that express *Piezo*2*, Calca*, or both. Data from N=20 images from 4 mice.

## Discussion

Bladder sensory neurons play a fundamental role orchestrating the voiding reflex and encoding visceral pain. While the existence of multiple subtypes of bladder-innervating DRG neurons has been recognized for a long time, progress in defining their role in normal and pathologic voiding as well as visceral pain has been hindered by the lack of pharmacological and genetic tools to target them. Here we used retrograde tracing and immuno-FISH to define molecular markers for the DRG neuron subtypes that innervate the urinary bladder. We initially evaluated the specificity of previously identified sensory neuron markers from available transcriptomic data in cross-section of unlabeled lumbosacral DRG [3–5, 20]. Of the selected markers, some labeled unique neuronal populations at the lumbosacral level (e.g., *Mgpra3*, *Mrgprd*, *Sst*), while others showed overlap with other markers to varying extent. Given that a single molecular marker may be insufficient to define a neuronal population, we employed a combinatorial approach integrating multiple markers and a binary (present/absent) expression scoring system to delineate the subtypes of DRG neurons that innervate the urinary bladder. Our studies indicate that the urinary bladder is innervated by at least four primary DRG neuron populations, three peptidergic that co-express *Calca*, and another non-peptidergic that expresses *Ntrk2* and *Piezo2* (Fig. 8). We anticipate that the markers identified in this study will facilitate selective access to specific populations with the arsenal of genetic tools currently available, including Cre and Flp mouse lines.

**Figure 8.**
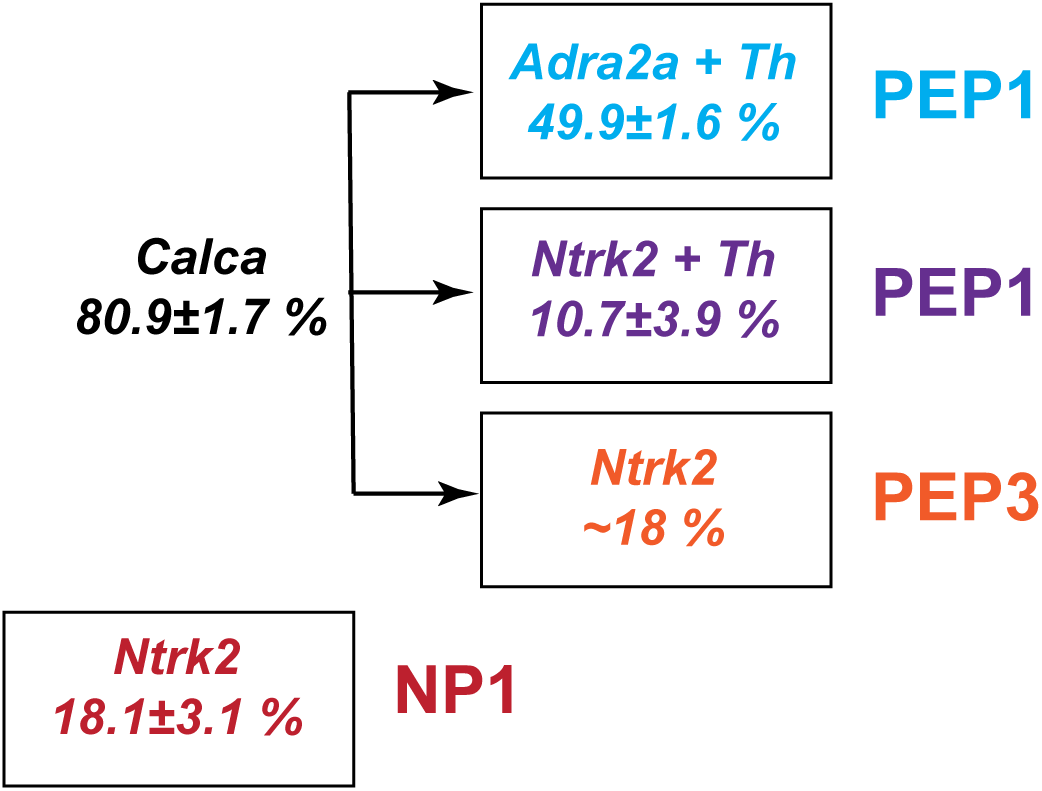
*Calca*, *Ntrk2*, *Th* and *Adra2a* delineate four major subtypes of bladder-innervating sensory neurons. Immuno-FISH was performed on retrogradely labeled bladder sensory neurons. The total population of DRG neurons that innervate the urinary bladder separates into four main non-overlapping classes: three subsets of peptidergic neurons that express Adra2a/*Th*, *Ntrk2*/*Th*, and *Ntrk2*, respectively, and a distinct non-peptidergic population that expresses *Ntrk2*. Percentage of bladder-innervating neurons in each group was calculated by averaging the results from the assays combining retrograde tracing and immuno-FISH. The percentage of *Calca*^+^/*Ntrk2*^+^ was calculated by subtracting the percentage of Adra2a^+^/*Th*^+^, *Ntrk2*^+^/*Th*^+^, and *Ntrk2*^+^ from the total population.

Retrograde tracing studies in rats revealed that roughly 5-6 % of DRG neurons at the lumbosacral level (i.e., L6-S2) project to the urinary bladder [21]. Given the low abundance of DRG neurons that innervate this organ, a pertinent question is whether they are among the molecular subtypes of DRG neuron clusters described by prior transcriptomic studies. We discovered that three peptidergic sensory neuron subtypes that innervate the urinary bladder are defined by combination of markers different to those described for general somatosensory or colon-innervating DRG neuron subtypes [7, 20], suggesting they encompass unique populations not previously identified in transcriptomic studies. For instance, we found that among the lumbosacral bladder-innervating neurons, almost all the *Adra2a*^+^ neurons express *Th*. On the contrary, of the whole population of lumbosacral *Adra2a*^+^ DRG neurons, only 34.2±9.3 % express *Th*, and of the *Th*^+^ neurons only 20.1±7.6 % express *Adra2a*. Moreover, the overlap detected between *Ntrk2* and *Th* within the whole population of bladder sensory neurons is negligible; nevertheless, a discrete subset, accounting for 10.7±3.9% of lumbosacral bladder-innervating neurons co-expresses both markers. Bladder-innervating DRG neurons are of low abundance and are distributed across a few axial levels of the spinal cord. As a result, unbiased clustering may not have been able to recognize them as separated entities. For instance, if 5% of the sensory neurons at that lumbosacral level (L6-S2) innervate the urinary bladder and 10% of them are Ntrk2^+^/Th^+^, then only 1 out of 200 neurons at this level will meet both criteria. Given that the studies with naive (non-bladder-traced) and retrogradely labeled DRG were done under identical conditions, we can reasonably conclude that the identified populations are genuinely distinct bladder-innervating neurons.

To the best of our knowledge, a combination of multi-labeling techniques and retrograde tracing, such as the used in this study, have not been applied to define sensory neuron populations. Spatial transcriptomics delivers high-resolution gene expression maps within the native architecture of tissue and can be combined with immunofluorescent staining, but the low abundance of bladder-projecting DRG neurons makes this approach impractical for the identification of neuronal populations and specific markers to label them. Immuno-FISH offers significant advantages over other technologies including simultaneous detection of transcripts and protein, preservation of spatial and anatomical context and high sensitivity for low abundance transcripts at single molecule level. Taken together, our study underscores the importance of validating the specificity of transcriptomically identified markers using immuno-FISH and highlights the need to define the DRG neuron populations that innervate specific tissues and organs using multiple approaches.

*Bmpr1b*, *Adra2a*, *Th* and *Ntrk2* have been previously shown to label distinct afferent clusters in the colon and skin, each with exclusive peripheral and central terminations as well as physiological properties [7, 20]. Our FISH studies of unlabeled lumbosacral DRG neurons revealed some overlap among particular combinations of these markers. An explanation for this discrepancy is that the unlabeled, double-positive neurons in our FISH study are not necessarily colon or skin afferents, and therefore express different combinations of markers. Alternatively, this discrepancy may be due to the different approaches used to identify neuronal populations: unbiased clustering of transcriptomic data, based on expression levels of multiple genes, versus immuno-FISH. Binary present/absent FISH scoring could be picking up low-level co-expression that a clustering algorithm would treat as negligible/noise and assign to one dominant class. Similarly, we found that a small percentage of *Adra2a*^+^ bladder-innervating neurons did not express TH when using an antibody to detect the protein, but most *Adra2a*^+^ neurons expressed *Th* when a probe is used to detect message. This could be explained by differences in the sensitivity of antibody- and probe-based detection systems toward their respective targets (protein versus mRNA). scRNA-seq and snRNA-seq allow for unbiased classification of neuron types [3–6], but identifying markers to target specific neuronal subsets remains a major challenge in neuroscience and other fields. Here we developed a simple approach for the characterization of bladder-innervating DRG populations that combines retrograde tracing, immuno-FISH and a present/absent scoring approach. The high sensitivity of immuno-FISH makes this approach well-suited for validating markers for morphological and functional studies. Using this approach, we identified four major sensory neuron subtypes that innervate the urinary bladder; three of them *Calca*^+^/*Adra2a*^+^/*Th*^+^, *Calca*^+^/*Th*^+^/*Ntrk2*^+^, or *Calca*^+^/*Ntrk2*^+^ have not been described previously in deep transcriptomic at studies. Thus, an important open question is whether the identified transcriptomic clusters represent the full complement of DRG neuronal populations, or whether additional and yet unidentified populations exist, for instance, those that innervate other visceral organs. Addressing this question will be important for building a complete map of the sensory neurons that innervate internal organs.

## Data availability

Raw data that support the findings of this study are available on request from the corresponding author.

## Author contributions

MDC and XS conceived the study. MDC and GMM performed retrograde tracing. XS performed FISH, immuno-FISH experiments including image capture and image analysis. MDC wrote the original draft and assembled the figures with input from all authors. All the authors approved the manuscript.

